# Temporal decoding of blood flow derived mechanical cues driving liver regeneration

**DOI:** 10.64898/2026.07.13.738320

**Authors:** Xinyu Shu, Guangyao Chen, Chaoyang Song, Yan Zhang, Shouqin Lü, Yu Du, Mian Long

**Author notes:** Correspondence (Y.D.), (M.L.).

## Abstract

Liver regeneration is initiated by rapid vascular changes, yet how blood flow-derived mechanical cues are decoded by liver sinusoidal endothelial cells (LSECs) remains unclear. Here, we found that partial hepatectomy generates temporally distinct mechanical cues *in vivo*, with a transient rise in shear stress followed by progressive sinusoidal dilation and endothelial stretch. To dissect these forces, we developed a liver regeneration chip that reconstructs sinusoidal architecture and enables independent or coupled manipulation of shear stress and mechanical stretch. Shear-dominant, stretch-dominant, and coupled mechanical modalities induce divergent LSEC regenerative programs involving extracellular matrix remodeling, cell-cycle regulation, cytoskeletal organization, and angiocrine signaling. Mechanistically, force-specific pathways, including Wnt, HIF-1, NF-κB, and Piezo1-associated signaling, mediate these outputs. Inhibition of these pathways after partial hepatectomy impairs hepatocyte proliferation and survival. These findings reveal that LSECs temporally decode blood flow-derived mechanical forces into distinct regenerative outputs, establishing endothelial mechanotransduction as an upstream regulator of liver regeneration.

**HIGHLIGHTS:**

■ Partial hepatectomy decouples transient shear from progressive stretch *in vivo*.
■ A liver regeneration chip recreates structure and distinct mechanical modalities in sinusoids.
■ Distinct mechanical modalities encode divergent LSEC regenerative programs.
■ Force-specific LSEC mechanotransduction supports hepatocyte proliferation and survival.

## INTRODUCTION

The liver has a remarkable capacity to restore mass and function after injury or surgical resection. Following partial hepatectomy, regeneration is initiated within minutes to hours prior to overt hepatocyte proliferation, indicating that early microenvironmental cues rapidly prime the remnant liver for growth^1,2^. Among those non-parenchymal cells that coordinate this response, liver sinusoidal endothelial cells (LSECs) occupy a central position at the interface between blood flow and hepatocytes^3–5^. By sensing changes in the vascular milieu and releasing angiocrine factors, LSECs regulate extracellular matrix remodeling, inflammatory and mitogenic signaling, and hepatocyte cell-cycle entry during the earliest phase of regeneration^2,5–9^.

Meanwhile, partial hepatectomy abruptly alters hepatic hemodynamics, exposing the remnant sinusoidal endothelium to increased blood flow, shear stress, and mechanical stretch^10–12^. Although hemodynamic changes have long been implicated in liver regeneration, how these blood flow-derived mechanical cues are mechanosensed by LSECs remains poorly understood^7,13,14^. In particular, shear stress and circumferential stretch are often considered together as a single mechanical consequence of increased flow, yet they represent distinct physical inputs that may differ in timing, cellular sensing mechanisms, and biological outputs^9,11^. Whether these mechanical modalities are temporally separated *in vivo* and how they differentially regulate LSEC angiocrine programs during regeneration initiation have not been directly evidenced.

A major barrier to address this question is the lack of experimental set-ups that preserve sinusoidal architecture while allowing controlled manipulation of mechanical forces. *In vivo* models retain physiological complexity but cannot decouple shear stress, pressure, and tissue deformation^15,16^. Conventional *in vitro* flow chambers and stretch devices isolate individual forces but impose different geometries, substrates, and stiffness, making direct comparison difficult^11,14^. Thus, a unified biomimetic platform is needed to dissect how LSECs decode blood flow-derived forces in a regeneration-relevant context^17^.

Here, we combine intravital imaging with a liver regeneration chip that reconstructs the sinusoidal structures and enables independent or coupled control of shear stress and stretch on endothelium. We show that partial hepatectomy generates temporally decoupled mechanical cues *in vivo*, with transient shear stress followed by progressive sinusoidal stretch. Using this chip, we demonstrate that these mechanical modalities drive distinct LSEC transcriptional and angiocrine programs linked to early liver regeneration, revealing a temporal decoding mechanism by which blood flow is converted into endothelial regenerative signals.

## EXPERIMENTAL MODEL AND STUDY PARTICIPANT DETAILS

### Isolation and culture of primary murine hepatic cells

Primary hepatic cells were isolated from 8- to 10-week-old male C57BL/6 mice (Vital River Laboratories, China). All animal experimental procedures were approved by the Animal and Medicine Ethical Committee of the Institute of Mechanics, Chinese Academy of Sciences. Liver homogenate was obtained by two-step collagenase digestion and filtered by a 70-μm cell strainer to remove the undigested fragments as previously described^18^. The collected cell suspension was then centrifuged at 54×*g* for 3 min at 4°C. The pellet was used for hepatocyte isolation and the supernatant was used for LSEC isolation.

For hepatocyte isolation, the pellet was resuspended in 36% Percoll (Sigma-Aldrich, USA) solution and centrifuged at 300×*g* for 5 min, then washed with calcium-free Hank’s balanced salt solution (HBSS, Macgene, China) again and resuspended in culture medium (high glucose DMEM supplemented with 10% FBS, 100 μg/ml streptomycin and 100 U/ml penicillin) at a final concentration of 0.3 million cells/ml and used immediately.

For LSEC isolation, the supernatant was centrifuged at 500 × *g* for 8 min. The pellet was resuspended with 3 ml of 24% Optiprep solution (Axis-Shield, Norway), then sequentially overlaid with 17.6% Optiprep, 11.2% Optiprep and 3 ml DMEM for density gradient centrifugation at 1400 ×*g* for 18 min. LSECs and Kupffer cells were obtained from the layer between 11.2% and the 17.6% Optiprep, followed by washing in HBSS and centrifugation at 1400 ×*g* for 8 min. The pellet was then resuspended with 90 μl DPBS and incubated with 10 μl anti-mouse CD146 (LSEC) MicroBeads (Miltenyi Biotec, Germany) at 4°C for 15 min, then centrifuged at 300×*g* for 10 min. The pellet was resuspended with 500 μl DPBS and applied to the MS column (Miltenyi Biotec). Unlabeled cells were washed through the column three times with 500 µL DPBS and discarded. The LSEC fraction was flushed out in 1 ml DPBS by firmly applying the plunger supplied with the column. The cell suspension was then centrifuged at 300×*g* for 10 min and resuspended in Endothelial Cell Medium (ECM, ScienCell, USA) at a final concentration of 2.5 million cells/ml and used immediately.

### 2/3 Partial hepatectomy model

2/3 partial hepatectomy based on the classic method by Higgins and Anderson^19,20^ was performed in 8-week-old male C57BL/6 mice as described. Briefly, a midline incision was created in the upper abdomen. The left and median lobes were ligated with a 5L0 silk suture (Shinva Medical Instrument Co., Ltd., China) and then removed. The laparotomy incision was closed with a running suture. For sham operation, mice received an identical incision and liver exposure. The liver lobes were gently mobilized with saline-moistened swabs, but no ligation or tissue removal was performed.

### Detection of RBC velocity and vessel diameter in sinusoids

Freshly isolated murine red blood cells (RBCs), labelled with Vybrant™ DiD cell-labeling solution (Thermo Fisher Scientific, USA), were administered intravenously *via* the tail vein together with Evans Blue (Solarbio, China). Liver exposure for intravital imaging was performed as described^21^. After midline laparotomy, the mouse was placed in right lateral decubitus. The caudate lobe was gently exteriorized and positioned directly on the optical window of a customLdesigned imaging stage. The lobe was stabilized with a Kimwipe (Kimberly-Clark, USA) and kept moist with DPBS throughout imaging. TimeLlapse imaging was performed on a spinning-disk confocal microscope equipped with an inverted microscopic platform (Olympus, Japan), a spinning-disk confocal scanner unit (Yokogawa, Japan), and an sCMOS camera (Photometrics, USA) with acquisition intervals at 10-20 ms for RBC tracking and 50 ms for diameter measurement. The collected images were processed using Imaris software (Oxford Instruments, UK) for cell tracking to calculate RBC velocities, and sinusoidal diameters were quantified using AngioTool software (National Cancer Institute, USA)^22^. Wall shear stress (τ) in hepatic sinusoids was estimated based on Poiseuille’s law as τ = 4η*v/r*, where η is the dynamic viscosity of blood, *v* is the mean red blood cell velocity, and *r* is the sinusoidal radius. A blood viscosity of 4 cP was used for the calculation^23^.

### Inhibitor treatment in 2/3 PHx model

Mice received intraperitoneal injections of the Wnt inhibitor WntLC59 (Selleck, USA; 20 mg/kg), HIF1 inhibitor PXL478 (MedChemExpress, USA; 50 mg/kg), NFLκB inhibitor PDTC (MedChemExpress, USA; 50 mg/kg), Piezo1 inhibitor GsMTx4 (MedChemExpress, USA; 2 mg/kg), or vehicle control, respectively, at 3 h before 2/3 PHx surgery. A second injection was administered 24 h after surgery and mice were sacrificed at 48-h postLhepatectomy with liver samples collected. Part of each liver sample was fixed, paraffinLembedded, sectioned, and processed for immunofluorescence staining. The remaining tissue was homogenized for RNA extraction and qPCR analysis.

## METHOD DETAILS

### Fabrication of the LRC

Liver Regeneration Chips (LRCs) were microfabricated by replica molding using 3D-printed masters. Briefly, the polydimethylsiloxane (PDMS, DowLCorning, USA) replicas were bonded to glass coverslips (Thermo Fisher Scientific, USA) and filled with collagen gels. The PDMS devices were incubated with 0.01% polyLLLlysine (Sigma-Aldrich, USA) solution for 1 h and 1% glutaraldehyde (Sigma-Aldrich) for 15 min to enhance collagen adhesion and then washed three times for 5 min each, ultrasonicated and soaked in water overnight. Steel acupuncture needles (Lejiu, China; 200 μm-diameter) were incubated in 1% BSA solution for 1 h and then inserted into the devices. All devices were sterilized under UV for 20 min. Rat tail type I collagen (Corning, USA) solutions at 2.5 or 7.8 mg/ml containing 1× DMEM medium, 10 mM HEPES, 0.1 M NaOH, and 0.035% NaHCO_3_ were infused into devices through the 2-mm ports and polymerized at 37°C for 20 min. DPBS was then added to all ports, and the needles were removed gently 12 h later. For LSEC seeding, cell suspension at 2.5 million cells/ml was introduced into the 4-mm reservoir ports. LSECs were allowed to adhere to the top surface of the channel for 5 min, and then the device was flipped for another 5 min as described^3^. After 1 h, nonadherent cells were rinsed away with ECM. Adherent LSECs in LRCs were cultured in incubator at 37°C with 5% CO_2_ for 24 h to form monolayers before use. For hepatocytes seeding, cell suspension at 0.3 million cells/ml was introduced into the 3-mm side ports. Culture medium was replaced 4 h later to remove nonadherent cells. Hepatocytes in LRCs were cultured over 12 h before use.

### Application of coupled or decoupled mechanical loading in LRC

Microfluidic flow controller OB1 MK3+ (Elveflow, France) was connected to computer, and the output pressure and flow rate were monitored by Elveflow Smart Interface (ESI) software (Elveflow, France). The control system was configured with an external vacuum pump and air pump to serve as power sources. HBSS from the reservoir was driven and monitored by ESI through the tubing to connect LRC. A Microfluidic Flow Sensor (Elveflow, France) integrated downstream of the reservoir enabled real-time feedback control.

To obtain varying degrees of PDMS deformation, LRCs for stretch-shear coupled and stretch alone loading modes were filled with collagen gel at 2.5 mg/ml which is soft and easily deformable, whereas a 7.8 mg/ml gel, being stiff and resistant to deformation, was used for shear alone mode. Under stretch-shear coupled and shear alone modes, precisely controlled flow was applied to the LRC channel with the outlet unobstructed and the effluent continuously aspirated by a vacuum-driven pipette. Under stretch alone mode, ESI-controlled pressure was applied with the channel outlet occluded.

### Immunofluorescence staining

Cells in LRCs were rinsed three times with DPBS and fixed with 4 % paraformaldehyde (Solarbio) for 15 min at room temperature. Fixed cells were washed again three times with PBS and permeabilized with 0.2% Triton XL100 (Solarbio) for 2 h, then blocked with 1% BSA (SigmaLAldrich) in PBS at 4°C overnight. Primary antibodies diluted in 1% BSA in DPBS were incubated overnight at 4L with rocking and then rinsed three times with DPBS followed by an overnight rinse. Secondary antibodies, 4′,6LdiamidinoL2Lphenylindole (DAPI, Solarbio) and phalloidin (Thermo Fisher Scientific) were diluted in 1% BSA and incubated overnight at 4L with rocking. Cells were rinsed three times, followed by an overnight rinse with rocking, and were visualized by LSM 880 confocal laser-scanning microscope (Zeiss, Germany).

Liver paraffin sections were deparaffinized, rehydrated, and subjected to antigen retrieval in EDTA buffer (pH 8.0). Antigen retrieval was performed at 100°C for 15 min for Ki67 staining and at 90°C for 30 min for TUNEL staining. Ki67 staining was performed using a tyramide signal amplification (TSA) system (Servicebio). After blocking and incubation with antiLKi67 antibody (Servicebio), sections were treated with HRPLconjugated secondary antibody (Servicebio), followed by signal detection with iF555-Tyramide (Servicebio). Apoptosis was assessed using a TUNEL assay. Sections were incubated with TUNEL reaction mixture (Servicebio) for 1 h at 37°C. Nuclei were counterstained with DAPI. WholeLslide images were acquired using a fluorescence microscope (Nikon, Japan) and a digital slide scanner (Servicebio).

### LSEC permeability measurement

To measure the permeability of the LSEC lumen in LRCs, fluorescein isothiocyanate-dextran at 70 kDa (FITC-dextran, SigmaLAldrich) in DPBS was introduced into the reservoir and the diffusion was imaged real-time using an LSM 880 confocal microscope (Zeiss) at 10× magnification. The diffusive permeability coefficient was determined by monitoring dextran flux in the extraluminal compartment and then fitting the resultant diffusion profiles to a dynamic mass conservation equation, following a previously described method^24^.

### Hepatocyte function assays

For glycogen storage analysis, hepatocytes were stained by Periodic Acid-Schiff (PAS) staining kit (Solarbio) and visualized by EVOS M7000 Imaging System (Thermo Fisher Scientific). To detect the transport function in hepatocytes, (5-and-6)-carboxy 2′,7′-dichlorofluorescein diacetate (CDFDA, Sigma-Aldrich) was used at a final concentration of 10 μM. Cells were incubated for 20 min at 37°C before observation with EVOS M7000 Imaging System.

### Quantitative real-time PCR (qPCR)

Total RNA was extracted from LSECs using a commercial RNeasy Micro Kit (QIAGEN, Germany), and the first-strand complementary DNA (cDNA) was synthesized using a ReverTra Ace-α reverse transcription kit (Toyobo, Japan) according to the manufacturer’s instructions. qPCR was performed using GoTaq® qPCR Master Mix (Promega, USA) and measured by a QuantStudio 7 Flex system (Thermo Fisher Scientific). Gene expression was normalized to endogenous *Gapdh* and calculated using the 2^-ΔΔCT^ method. All primers used in this work were listed in Supplementary Table 1.

### Visualization of LSEC lumen diameter

LRC connected to the loading system was positioned on the stage of IX81 automatic inverted microscope (Olympus, Japan). Time-lapse differential interference contrast (DIC) images of LSEC lumens in LRC were captured by an electron-multiplying charge-coupled device camera (Andor, UK) at 0.5 s intervals for 5 min. Collected image sequences were analyzed by Fiji software ^25^. LSEC lumen strain was defined as (*D* − *D*L)/*D*L, where *D* is the lumen diameter under loading and *D*L is the initial lumen diameter without loading.

### Particle image velocimetry

To identify flow velocity in LSEC lumen of LRC, 2 μm FITC-labeled beads in HBSS were continuously perfused into LRC under coupled and decoupled loading modalities. Fluorescent beads in LSEC lumen were excited at 488 nm on IX81 automatic inverted microscope and their trajectories were captured by Andor camera with an exposure time of 40 ms per frame. Fluorescent trail lengths of beads were measured by Fiji software and their velocities were calculated.

### Atomic force microscopy

A JPK NanoWizard 4XP atomic force microscope (AFM) (Bruker, USA) was applied to measure the Young’s modulus of collagen gels at concentrations of 2.5 and 7.8 mg/ml. Briefly, force curves were obtained with a MLCT-C cantilever at approach and retraction velocities of 300Lnm/s with a loading force of 300 pN. The ZLlength was set to 1.0 µm. Force maps of 3L×L3Lµm areas with 8L×L8 sampling points were collected in three independent repeats. ForceLindentation curves were analyzed using the Sneddon model in JPKSPM Data Processing software (Bruker), assuming a conical tip with a halfLcone angle of 15° and a Poisson’s ratio of 0.5. Average Young’s modulus was calculated by the mean of the values at each point in the field-of-views (FOVs).

### RNA-sequencing analysis

Total RNA was extracted from LSECs obtained from the collagen gel after mechanical loadings as described (Fig. S3a) and each group was prepared with more than three biological replicates. RNA collected was sent to BGI Genomics (Shenzhen, China) for library construction and sequencing were performed on the DNBSEQ platform. Differential expression analysis was conducted using the DEGseq algorithm *via* the online Dr. Tom system developed by BGI, with |fold change| ≥ 1.5 and *q* value ≤ 0.05^26^. KEGG pathway enrichment analysis was performed on the upregulated differentially expressed genes on Dr. Tom system, and the top 25 enriched pathways were exported for visualization.

PreLranked gene set enrichment analysis (GSEA) was performed using the fgsea R package. Gene expression data were transformed to logL(TPML+L1) prior to analysis. For each comparison, genes were ranked by the logL fold change (logLFC) calculated as the difference between the mean expression of each mechanical modality and static control. Gene sets of interest comprised cell cycle (defined as the union of HALLMARK_G2M_CHECKPOINT and HALLMARK_E2F_TARGETS), cellular senescence (GO:0090398), KEGG regulation of actin cytoskeleton (mmu04810), KEGG cell adhesion molecules (mmu04514), inflammatory response (HALLMARK_INFLAMMATORY_RESPONSE), MHC class I antigen presentation (GO:0002474), and apoptosis (HALLMARK_APOPTOSIS). Only genes present in the expression matrix were retained. Enrichment was assessed using the fgseaMultilevel function. Adjusted *p*Lvalue < 0.05 was considered significant, and the normalized enrichment score (NES) was reported for each pathway.

Weighted gene coLexpression network analysis (WGCNA) was performed using the WGCNA R package^27^. A signed network was constructed with a softLthresholding power of 9. Modules were identified with a minimum size of 30 genes and merged at a cut height of 0.25. Module eigengenes were correlated with sample traits including stretch-shear, stretch and shear modalities. Modules with |correlation| > 0.5 and *p* < 0.05 were considered nominally significant, and their member genes were extracted for downstream analysis.

### Quantification and Statistical Analyses

Statistical significance was assessed using the Student’s *t*-test with equal variance if they passed the normality test (Shapiro-Wilk) or the Mann-Whitney test if not for two groups, and using one-way ANOVA test if they passed the normality test or ANOVA on RANKs if not for multiple groups. *p* < 0.05 was regarded as statistically significant and calculated with Prism 7 (GraphPad Software, La Jolla, CA, USA) and SigmaPlot 12.5 (San Jose, CA, USA). All data are presented as the mean ± standard error of the mean (SEM). Sample size was indicated in the figure legends.

### Data Availability

All data needed to evaluate the conclusions are presented in the text and/or the Supplementary Materials. Additional data related to this paper may be requested from the authors.

## RESULTS

### Partial hepatectomy generates temporally decoupled shear stress and stretch *in vivo*

To define the blood flow-derived mechanical cues experienced by LSECs during the initiation of liver regeneration, we performed intravital imaging of liver sinusoids in mice subjected to two-thirds partial hepatectomy (PHx)^28,29^. Using fluorescently labeled liver sinusoids and RBCs, sinusoidal blood flow velocity and vessel diameter were quantified in the remnant liver (caudate lobe) at 1 to 5 h and day 3 after surgery, allowing dynamic estimation of shear stress and endothelial stretch during the earliest regenerative phase (Fig. 1a-b, S1a-b). Consistent with acute hyperperfusion of the remnant liver, PHx induced an immediate increase in sinusoidal blood flow velocity which peaked at 10.3 dyn/cm^2^ within the first hour and subsequently declined to preoperative levels of 7.5 dyn/cm^2^ by 5 h post resection (Fig. 1b). In contrast, sinusoidal diameter exhibited a distinct temporal pattern. Rather than showing an acute increase, vessel diameter expanded progressively throughout the 5-h observation period after PHx (Fig. 1b), indicating endothelial stretch increased continuously over time and remained elevated even after shear stress had largely returned toward baseline levels.

**Figure 1.**
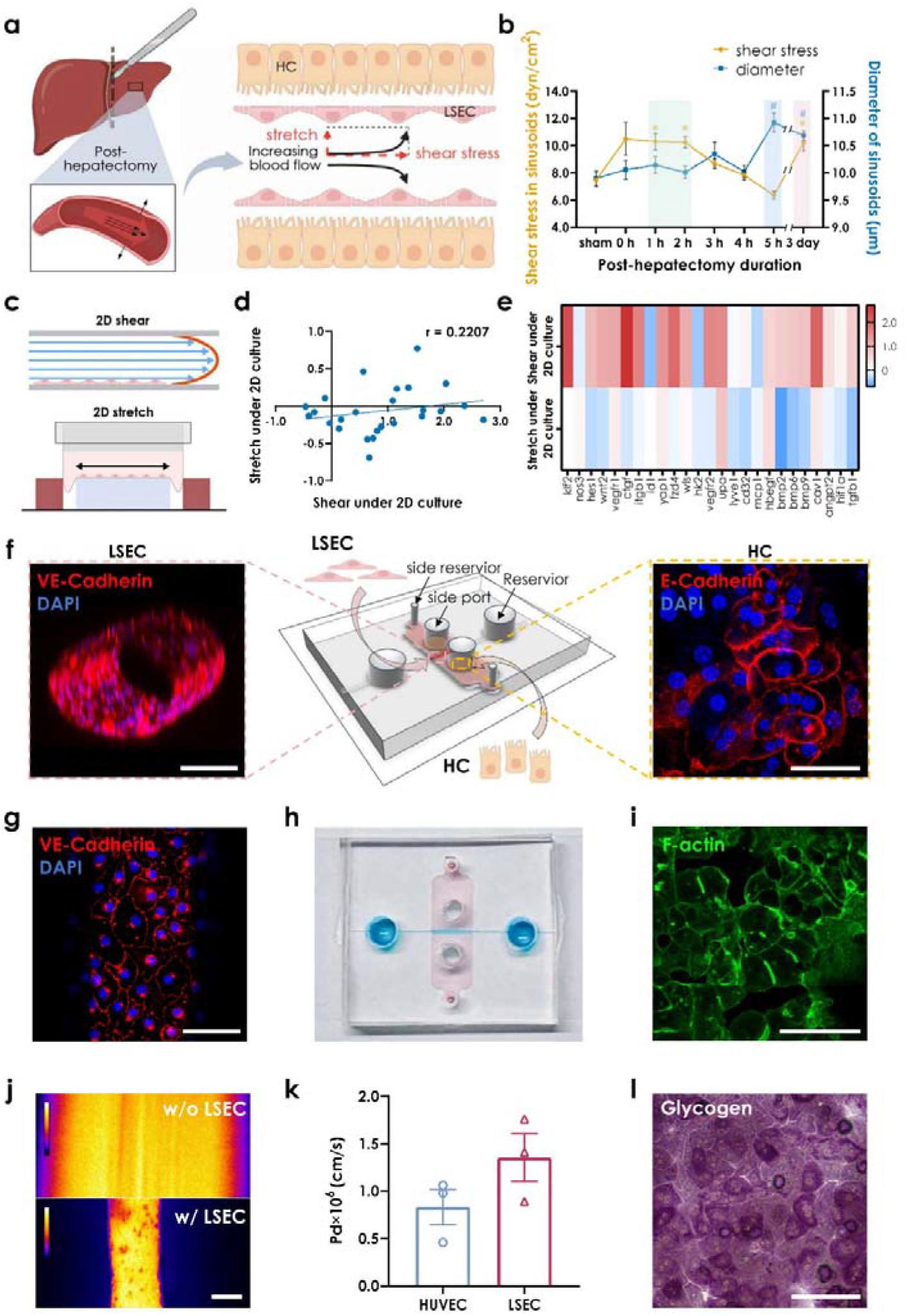
Fabrication and characterization of a co-culture microfluidic platform for liver regeneration. *a*, Schematic of hemodynamic alterations in hepatic sinusoids after partial hepatectomy. *b*, Post-hepatectomy profiles of sinusoidal shear stress and diameter variations over time. All data were presented as the mean ± SEM, *n* = 3-4 mice for each time point. The semiLtransparent regions indicated temporal phases dominated by shear (*green*), stretch (*blue*) and stretch-shear (*red*) loading, respectively. *c*, Conventional 2D culture and loading devices for shear or stretch alone. *d*, *e*, Pearson correlation coefficient (*d*) and expression heatmap (*e*) of liver regeneration-associated genes from 2D shear or stretch alone. Positive or negative values indicated the up- or down-regulated genes under each mechanical stimulus. Data were obtained from previously published data of LSECs under 2D stretch (Wu et al., 2024) and from LSECs under 2D shear in this study. *f*, Construction of liver regeneration chip (LRC). Isolated primary murine LSECs were seeded in collagen gel scaffolds with primary murine HCs arranged in side ports (*middle*). LRC architecture was presented by 3D fluorescence reconstruction, stained for VE-Cadherin (*red*) and DAPI (*blue*) for LSECs in lumen (*left*) and for E-Cadherin (*red*) and DAPI (*blue*) for HCs in side ports (*right*). Scale bar = 100 μm (left) and 50 μm (right). *g*, Representative 2D prejected immunofluorescence image of LSECs in channel were stained for VE-Cadherin (*red*) and DAPI (*blue*). Scale bar = 50 μm. *h*, Optical image of LRC device. *i*, Representative immunofluorescence image of bile canaliculi between HCs were stained for F-actin (*green*). Scale bar = 100 μm. *j*, Heatmap of fluorescence intensity of 70-kDa FITC-labelled dextran in LRC channel without (*upper*) or with LSECs (*lower*). *k*, Quantification of diffusive permeability (Pd) of 70-kDa dextran across endothelial barrier of HUVEC and LSEC lumen. All data were presented as the mean ± SEM, *n* = 3 independent repeats. *l*, Assessment of hepatocyte glycogen storage capacity by Periodic Acid-Schiff stain (*purple*). Scale bar = 125 μm.

These findings reveal that blood flow-derived mechanical forces are not transmitted to the regenerating liver as a single mechanical input. Instead, PHx generates two temporally distinct mechanical modalities: a transient surge in shear stress and a progressively accumulating stretch signal. This temporal separation suggested that LSECs are exposed to sequential, rather than synchronous, mechanical cues during regeneration initiation, potentiating that distinct mechanical modalities may differentially regulate endothelial regenerative programs.

### A liver regeneration chip reconstructs sinusoidal niche and enables force-specific interrogation

To investigate how temporally distinct mechanical cues regulate liver regeneration, we sought to build a minimal sinusoidal model that preserves the key architectural features required for LSEC mechanotransduction while enabling independent or coupled control of shear stress and mechanical stretch. We first assessed whether conventional *in vitro* platforms allow direct comparison of shear- and stretch-induced responses in LSECs^11,14,16,30^, when transcriptomic profiling of primary LSECs were exposed to laminar shear stress using a parallel-plate flow chamber or to cyclic stretch using a commercial FlexCell tension device (Fig. 1c). Comparison of differentially expressed genes (DEGs) revealed substantial divergence between the two modes and Pearson correlation analysis demonstrated low concordance between shear- and stretch-responsive transcriptional programs (Fig. 1d-e). These results suggested that platform-dependent differences in geometry, substrate mechanics, and parameter settings might undermine the comparison of force modalities, highlighting the need for a unified experimental system.

To address this limitation, we developed a liver regeneration chip (LRC) that integrates microfluidic technology with a pressure-driven pumping system to reconstructs key structural configuration and mechanical microenvironment of the liver sinusoid (Fig. 1f, h, S1c-f). The device consisted of a deformable endothelial channel embedded within a collagen matrix and flanked by hepatocyte compartments, thereby recapitulating the spatial organization of the sinusoidal niche (Fig. S1f). Primary LSECs formed a confluent tubular monolayer with intact VE-cadherin expressing cell–cell junctions and low permeability to fluorescent dextran, indicating the functional barrier of preserved endothelium (Fig. 1f-g, j-k, S1e)^24^. Meanwhile, primary murine hepatocytes established polarized epithelial monolayer (Fig. 1f, S1e) with well-defined bile canaliculi and glycogen storage capacity (Fig. 1i, l and S1f), indicating the preservation of essential hepatic functions. Collectively, these results demonstrated that the LRC reconstructs key sinusoidal features relevant to endothelial mechanotransduction during regeneration initiation, providing a biomimetic platform for understanding endothelial–hepatocyte interactions during liver regeneration.

A defining feature of this LRC is the deformable endothelial channel, which allows mechanical forces to be manipulated within the same sinusoidal architecture by dynamic modulation of both luminal flow and channel geometry in response to externally applied pressure in the stiffness-tunable gels (Fig. 2a, S2a). To determine whether this design enables force-specific mechanical interrogation, we systematically characterized shear stress and mechanical stretch under different loading patterns. Increased flow rate at constant intraluminal pressure selectively elevated shear stress with minimal changes in channel diameter, thereby generating a shear-dominant mechanical modality (Fig. 2b-e and S2c). In contrast, increasing pressure at constant flow rate induced progressive channel dilation with limited changes in flow velocities, generating a stretch-dominant mechanical modality (Fig. 2b-c, f-g). Simultaneous modulation of flow rate and pressure produced coordinated increases in stretch and shear, mimicking the coupled mechanical modality observed *in vivo* after partial hepatectomy (Fig. 1b). Quantitative analysis confirmed predictable control of force magnitude and channel deformation across all loading conditions (Fig. S2b-c). Together, these results establish the LRC as a biomimetic platform that not only reconstructs the structural and functional niche of the liver sinusoid but also enables force-specific interrogation of endothelial responses under independently controlled or physiologically coupled mechanical loading conditions.

**Figure 2.**
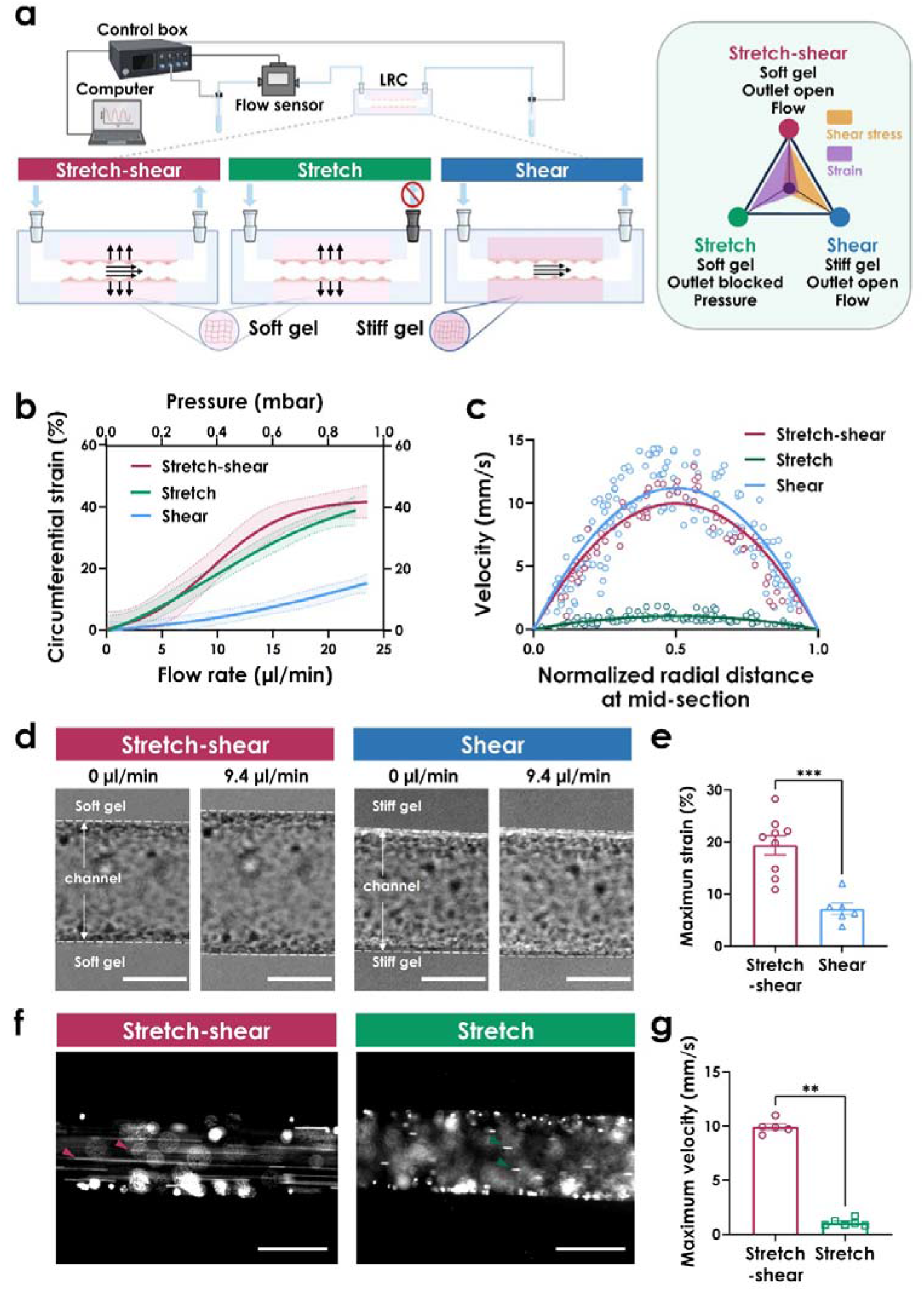
Implementation and validation of decoupled and coupled loading modes in LRC. *a*, Schematic of three mechanical loading modes: stretch-shear coupled, stretch or shear alone with respective mechanical cues indicated (*right*). *b*, LSEC lumen circumferential strain in LRC versus increasing flow loading or increasing pressure. *Solid lines* and *shadow regimes* indicated the measured data (*trend lines*) and the 95% confidential intervals, respectively. *c*, Distribution of fluorescent particles velocity in LSEC lumen along normalized radial distance of LRC at the mid-section (*points*). *Solid lines* were obtained by fitting the data with a centered second-order polynomial equation *y* = *a*·*x* - *b*·*x*² where *a* and *b* are fitting constants. *d*, *e*, Representative images of LSEC lumens under stretch-shear (*red*) and shear (*blue*) modes at null loading (*left*) or a typical flow rate of 9.4 μl/ml (*right*) (*d*), with the calculated maximum circumferential strain of LSEC lumens (*e*). *f*, *g*, Representative images of fluorescent particle tracking under stretch-shear (*red*; at a flow rate of 9.4 ml/ml) and stretch (*green*; at a pressure 0.4 mbar) modes (*f*), with the calculated maximum velocity in LRC lumens (*g*). *Red* and *green* arrows in *f* marked the motion trajectories of fluorescent particles during stretch-shear and stretch modes, respectively. Scale bar in *d* = 100 μm and *f* = 200 μm. All data in *e* and *g* were presented as the mean ± SEM, *n* = 3-4 independent repeats.

### Distinct mechanical modalities encode divergent regenerative programs in LSECs

Having established the ability to generate defined mechanical modalities in the LRC, we next sought to investigate whether shear-dominant, stretch-dominant, and coupled shear–stretch cues are decoded by LSECs into distinct functional programs. We performed RNA sequencing of LSECs exposed to the three mechanical modalities in the LRC, together with static controls (Fig. S3a). Pairwise comparison of global expression changes and Spearman correlations revealed the low to moderate relevance among three loading conditions, indicating that each modality elicited a distinct transcriptional response rather than different magnitudes of a common mechanical response (Fig. 3a). We then used gene set enrichment analysis (GSEA) to determine how these modality-specific transcriptional responses are mapped onto LSEC functions relevant to regeneration. Coupled stretch-shear loading significantly enriched those programs associated with cell cycle, apoptosis, actin regulation, and classical inflammatory response pathways (FDR < 0.05, NES > 1). In contrast, stretch -dominant loading downregulated cell adhesion molecules and immune-related gene sets (FDR < 0.05, NES < −1.7), whereas shear-dominant loading upregulated cellular senescence and MHC I antigen presentation pathways (FDR < 0.05, NES > 1.4) but downregulated classical inflammatory response pathways (FDR < 0.05, NES < −1.7) (Fig. 3b, S3b). These results suggested that distinct force modalities encode separable endothelial programs involved in proliferative activation, structural remodeling, immune regulation, and stress adaptation.

**Figure 3.**
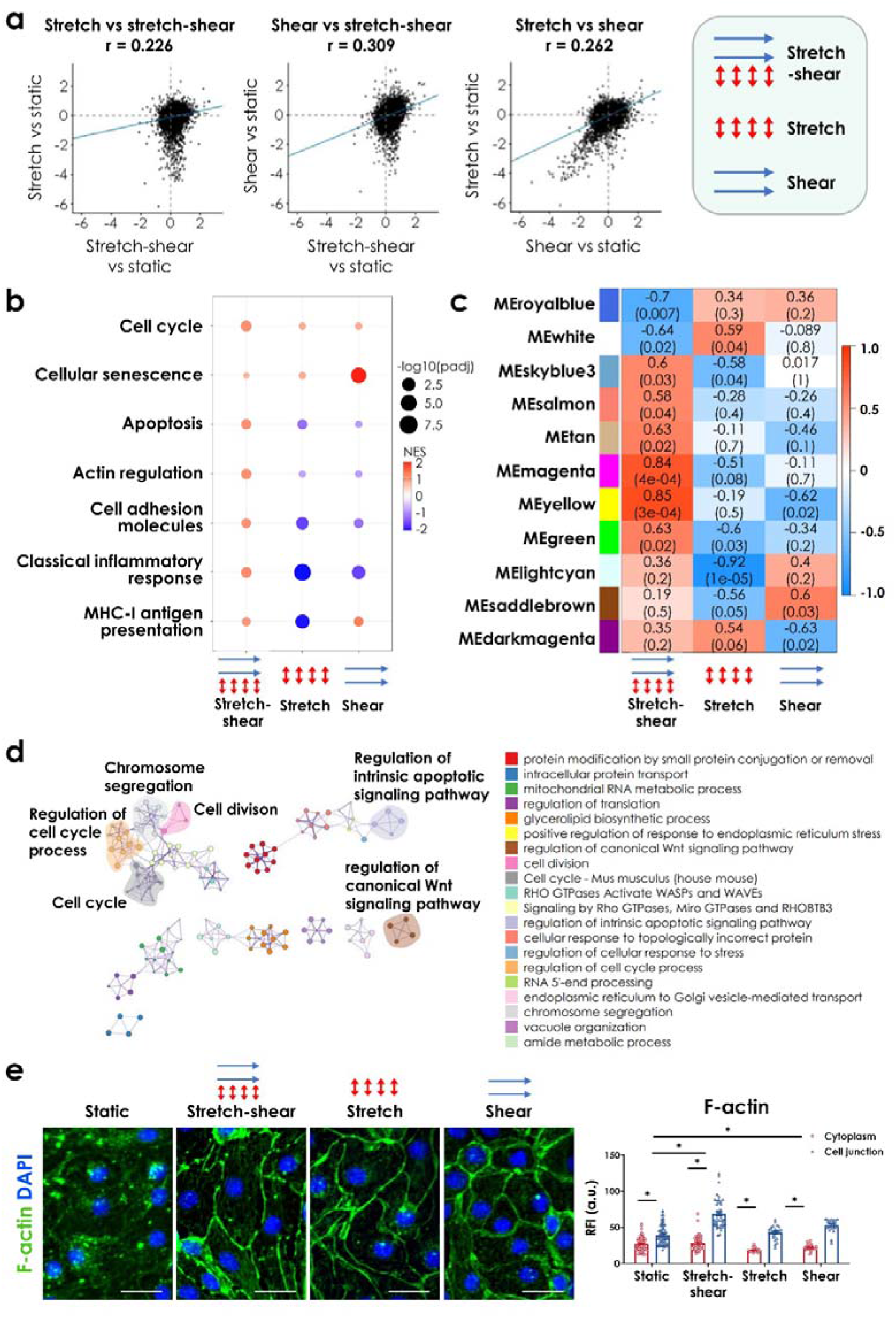
Transcriptional and cytoskeletal responses of LESCs to three mechanical modalities. *a*, Scatter plots with Spearman correlation coefficients (*r*) illustrated pairwise comparisons of the transcriptional expression changes among stretch-shear-coupled (input flow rate at 9.4 μl/min with estimated shear stress at 1.8 dyn/cm^2^ and circumferential strain at 20%), shear-dominant (input rate at 9.4 μl/min with estimated stress at 2.0 dyn/cm^2^ and strain at 3%) and stretch-dominant (input pressure at 0.4 mbar with estimated stress at 0.2 dyn/cm^2^ and strain at 20%) modes (relative to static). *b*, GSEA analysis of pathways associated with LSEC phenotype and function under three modes. *c*, WGCNALderived moduleLtrait correlation heatmap for significant modules under static and three loading modes. *d*, Network profile of subset of Gene Expression Omnibus (GEOs) derived from most significantly upLregulated module identified by WGCNA. GEO was represented by cluster identity and each term was represented as circle node visualized on Metascape. *e,* Representative fluorescent images of F-actin (*left*) and the quantified F-actin (*right*) distribution for cytoplasm and cell junction. Scale bar in *e* = 20 μm. All data in *e* were presented as the mean ± SEM, *n* = 3 independent repeats.

To further identify coordinated gene networks associated with each mechanical modality, we performed weighted gene co-expression network analysis (WGCNA). Among the modality-associated modules, the MEyellow module showed the most pronounced alteration and was specifically upregulated under stretch-shear modality. Metascape analysis of genes in the MEyellow confirmed the enrichment of cell cycle and apoptosis pathways as shown in the GSEA’s readouts (Fig 3.c-d), supporting the speculation that coupled mechanical cues induce an integrated regenerative state in LSECs rather than a simple additive response to shear or stretch cue alone. Noting that endothelial adaptation to mechanical stimulation requires coordinated remodeling of cytoskeleton and cell–cell junctions, we next examined the spatial organization of F-actin and VE-cadherin in LSECs under different mechanical modalities. Immunostainings revealed that stretch-shear loading enhanced actin enrichment at cell junctions whereas stretch alone reduced junctional VE-cadherin localization (Fig. 3e, S3c-d), confirming the differential endothelial responses under distinct modalities.

To further relate these LRC-induced programs to endothelial states during liver regeneration *in vivo*, we compared the mechanical modality-associated signatures with published scRNA-seq profiles of LSECs at day 3 postLPHx (Fig. S3f). Notably, the coupled shear–stretch signature correlated positively with LSECs at day 3 post-PHx, consistent with the coupled elevation of shear stress and stretch observed *in vivo* at this stage (Fig. 1b and Fig. S3f). Together, these results demonstrated that distinct blood flow-derived mechanical modalities encode divergent LSEC regenerative programs, with stretch-shear loading considerably recapitulating the *in vivo* regenerative signature.

### Force-specific mechanotransduction pathways mediate the regenerative effects of shear and stretch

Given that inhibition of mechanosensitive pathways in the liver has been shown to delay liver development and regeneration^4,7,11,31–33^, we sought to identify which mechanosensitive pathways are engaged by distinct mechanical modalities and whether they represent potential regenerative targets. Here KEGG enrichment was performed and the top 25 pathways revealed the stretch-shear specific activation of Wnt signaling, stretch specific activation of HIF-1 signaling, and shear specific activation of NF-κB signaling (Fig. 4a, S4a). In parallel, shear-induced changes in *Klf2* and its downstream targets mirrored the transcriptional profile of LSECs at early time after PHx, implicating Piezo1 as a potential upstream mechanosensitive regulator (Fig. S3e)^34^. To test whether these pathways contribute to mechanically induced angiocrine signaling, we pharmacologically inhibited those corresponding pathway candidates in LSECs exposed to defined mechanical modalities in the LRC. Inhibition of Wnt, HIF-1 or NF-κB signaling attenuated the regeneration-associated paracrine factors *Hbegf* and *Hgf*, together with upstream markers linked to LSEC-mediated regeneration (Fig. S4a). These results indicated that force-specific signaling pathways are required, at least in part, for the mechanoregulated angiocrine response of LSECs.

**Figure 4.**
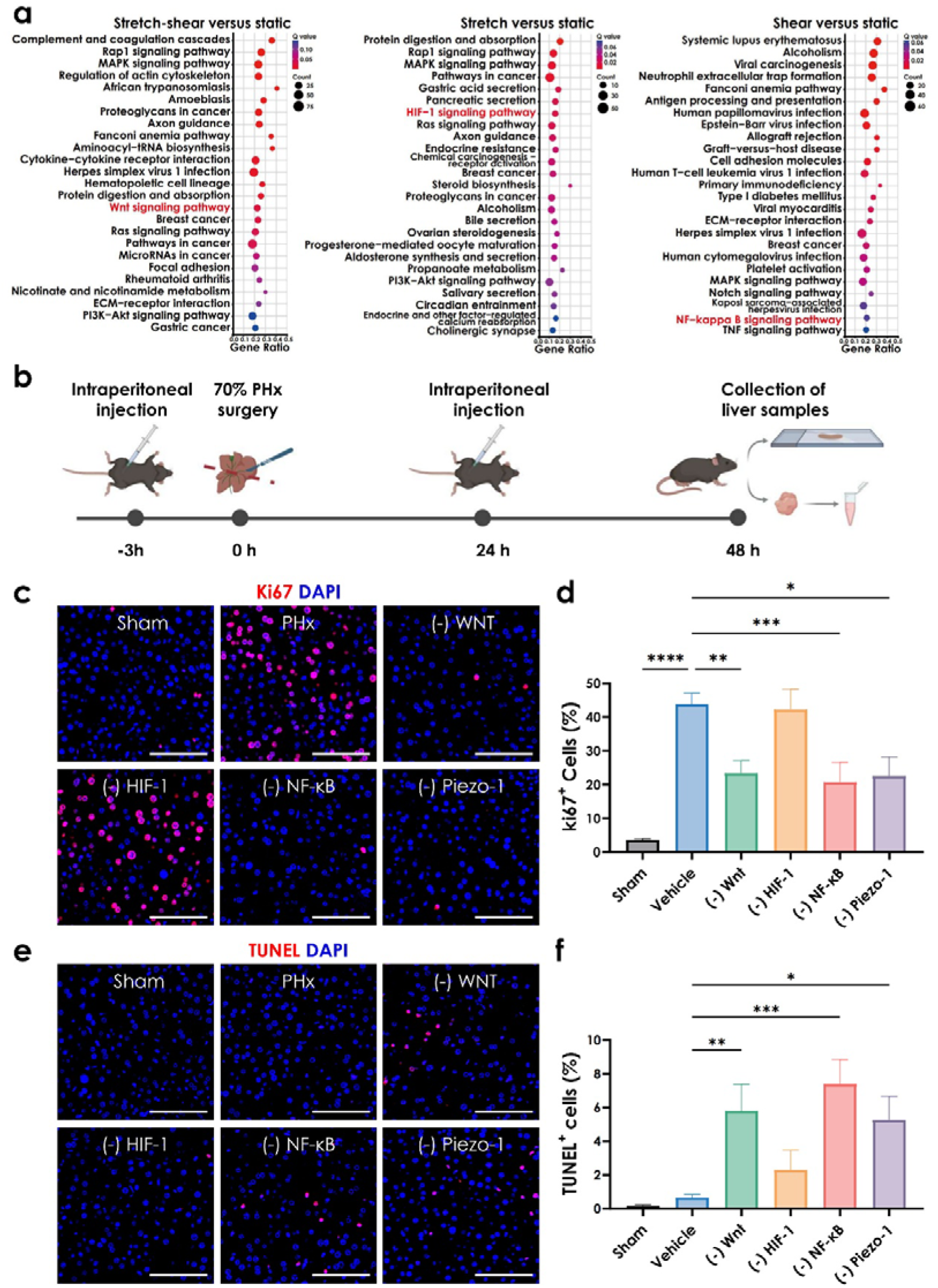
Validation of potent mechanosensitive targets *in vivo*. *a*, Bubble plots of KEGG pathway enrichment under three loading modes. *b*, Schematic of experimental setup to drug administration, 70% PHx surgery and sample collection in mice, created with Biorender. *c*, *d*, *e*, *f*, Representative fluorescence images of Ki-67 (*c*) and TUNEL (*e*)-positive hepatocytes in 70% PHx mice at 48 h post-treatment with different inhibitors (against Wnt, Piezo-1, NF-κB, or HIF1), together with the quantified percentages of Ki-67 (*d*) and TUNE- positive (*f*) hepatocytes. Scale bar in *c* and *d* = 100 μm. All data in *d* and *f* were presented as the mean ± SEM, *n* = 3-4 independent repeats.

We then investigated whether these mechanotransduction pathways influence liver regeneration *in vivo*. Mice were treated with inhibitors targeting Wnt, HIF-1, NF-κB, or Piezo1 signaling and subjected to two-thirds PHx, followed by analysis at 48 h post-surgery (Fig. 4b). Inhibitor treatment did not affect post-hepatectomy survival (Fig. S4b) and liver/body weight ratios were not significantly altered at 48 h post-PHx (Fig. S4c). However, inhibition of Wnt, NF-κB, or Piezo1 significantly reduced the proportion of Ki67-positive hepatocytes compared with vehicle-treated controls, consistent with reduced expression of proliferation-associated genes (Fig. 4c-d, S4d). Notably, TUNEL staining revealed increased apoptotic cells under same inhibition conditions (Fig. 4 e-f), indicating the impaired regenerative cell-cycle entry together with reduced survival support. Together, these results indicated that force-specific mechanotransduction in LSECs converts blood flow-derived mechanical cues into pro-regenerative signals that support hepatocyte cell-cycle entry and survival after PHx, highlighting this mechanical decoding process as a potential target for modulating liver regeneration.

## DISCUSSION

In this study, we identify a temporally orchestrated mechanical program that operates during the initiation of liver regeneration. By combining intravital imaging with a liver regeneration chip that enables controlled mechanical stimuli of LSECs, we show that partial hepatectomy does not simply expose the remnant liver to a uniform increase in mechanical force. Instead, blood-flow redistribution is translated into distinct mechanical modalities with different temporal profiles: shear stress rises rapidly and subsequently normalizes, whereas sinusoidal dilation and endothelial stretch progressively increase over the early regenerative window. This temporal decoupling provides a framework in which blood flow redistribution is not a uniform mechanical perturbation, but a sequence of force modalities that can be differentially interpreted by LSECs to initiate liver regeneration.

A central implication of our findings is that shear stress and mechanical stretch on endothelium should not be viewed as interchangeable consequences of increased flow. Rather, they appear to represent functionally distinct mechanical inputs that are differentially decoded by LSECs. The rapid shear response may act as an early sensing or triggering signals, alerting the sinusoidal endothelium to varied hemodynamic load immediately after tissue loss. In contrast, the gradual accumulation of stretch may provide a more sustained mechanical context that reinforces endothelial remodeling and angiocrine signaling. This division of labor may help explain a long-standing paradox in liver regeneration: although blood flow-derived shear stress can normalize within hours, the regenerative program continues to progress toward hepatocyte priming and proliferation^35^.

Methodologically, the LRC developed here was not intended to reproduce the full complexity of liver regeneration, but to be viewed as a controlled system between *in vivo* observation and mechanistic perturbation. Its value lies in reconstructing the minimal sinusoidal architecture required to isolate how LSECs respond to defined shear and stretch cues within the same microenvironment, which meets recent research trend by enabling *in vitro* investigation of biophysical cues in disease process^36^. Conventional flow chambers and stretch devices are useful for isolating single inputs, but they impose different geometries, substrate stiffnesses and mechanical parameters, making it hard to share direct comparisons between shear-and stretch-induced responses. By characterizing the temporal dynamics of mechanical cues *in vivo*, decoupling its component modalities in the LRC, and testing signal pathway relevance after PHx, our study uses this LRC chip to convert a complex hemodynamic response into experimentally tractable mechanical states.

Our transcriptomic analyses further indicate that distinct mechanical modalities induce highly divergent LSEC states rather than merely different magnitudes of a common response. Mechanically-regulated programs are associated with ECM remodeling, cell cycle, apoptosis, senescence, and actin organization, suggesting that LSECs integrate physical signals into coordinated angiocrine and structural responses. These programs are closely aligned with early events in regeneration, including matrix remodeling, release or presentation of pro-regenerative cues, endothelial adaptation, and hepatocyte priming^2^. Thus, endothelial mechanotransduction may function not only as a sensor of altered blood flow but also as an organizer of the early regenerative niche.

This work also has broader implications for regenerative medicine. Current strategies to accelerate liver regeneration often focus on growth factors, cytokines, or cell-based interventions^12,37^. Our findings suggest that the physical microenvironment itself may provide an upstream regulatory layer that shapes endothelial regenerative outputs. The identification of force-associated Wnt, HIF-1, NF-κB, and Piezo1-related pathways further suggests that temporal mechanical decoding is linked to the broader signaling network that regulates liver regeneration. These pathways should not be interpreted as a simple list of therapeutic targets, but rather as candidate nodes through which LSECs connect blood flow-derived mechanical inputs to established regenerative programs. It is also noted that our inhibitor experiments establish functional requirement for these pathways after PHx, but do not determine whether this pathway activation is sufficient to enhance regeneration or whether their effects are strictly LSEC-specific *in vivo*. Future studies using endothelial-specific perturbation, pathway reporters, and gain-of-function approaches will be needed to define how each pathway contributes to LSEC angiocrine output and whether mechanical decoding can be modulated to promote liver repair without driving excessive or pathological growth.

Another limitation is that the mechanical decoupling achieved in the LRC is experimentally defined rather than physically complete. Current mechanical modalities didn’t investigate intraluminal pressure, matrix stiffness and geometry, which may also contribute to endothelial responses. These conditions should therefore be interpreted as force-biased mechanical modalities rather than fully isolated physical variables. Moreover, the current model does not include Kupffer cells, hepatic stellate cells, infiltrating immune cells, or whole-organ metabolic and neurohumoral regulation, all of which may modulate LSEC mechanotransduction *in vivo*^38^. Future efforts to make the model physiologically relevant should also incorporate other cell components that may play a role in liver regeneration.

Finally, our study focuses on the initiation phase of liver regeneration and therefore does not address the role of mechanical decoding during later phases of liver growth, or termination of regeneration. This distinction is important because the restoration of hemodynamic changes overlaps with the termination of liver regeneration which prevents excessive or maladaptive growth^1,7^. It also remains unknown whether the mechanical decoding mechanism is preserved in diseased livers, where the fibrosis, steatosis and portal hypertension may reshape force transmission and LSEC mechanosensitivity. It will also be important to validate these mechanisms in human liver tissues and disease contexts, which may alter the mechanical decoding capacity of LSECs^39^. Overall, our study provides proof-of-concept evidence that temporal decoding of blood flow-derived mechanical cues represents a mechanobiological mechanism regulating the earliest phase of liver regeneration.

## ACKNOWLEDGMENTS

We thank Weiguo Fan and Yali Li (Center for Excellence in Molecular Cell Science, Shanghai Institute of Biochemistry and Cell Biology, University of Chinese Academy of Sciences, Chinese Academy of Sciences) for their technical support in hepatectomy surgery and intravital microscopy. This work was supported by National Natural Science Foundation of China grants T2394514, T2394510, 32250017, 32130061, and 12372320.

## AUTHOR CONTRIBUTIONS

M.L., Y.D. and X.Y.S. conceptualize the idea of the project; X.Y.S. and Y.D. performed the experiments and acquired the data with help from G.Y.C.; Y.D. andC.Y.S. designed the device mold. X.Y.S., Y.D. and M.L. analyzed the data; X.Y.S.,Y.D. and M.L. wrote the manuscript.

## Data and code availability

All sequencing data have been deposited at ScienceDB repository and are available under the following URL: https://www.scidb.cn/anonymous/ZnFBZmlt. DOI is https://doi.org/10.57760/sciencedb.42501. The data that support the findings of this study are available from the corresponding author upon reasonable request.

---

**Supplementary figure 1.**
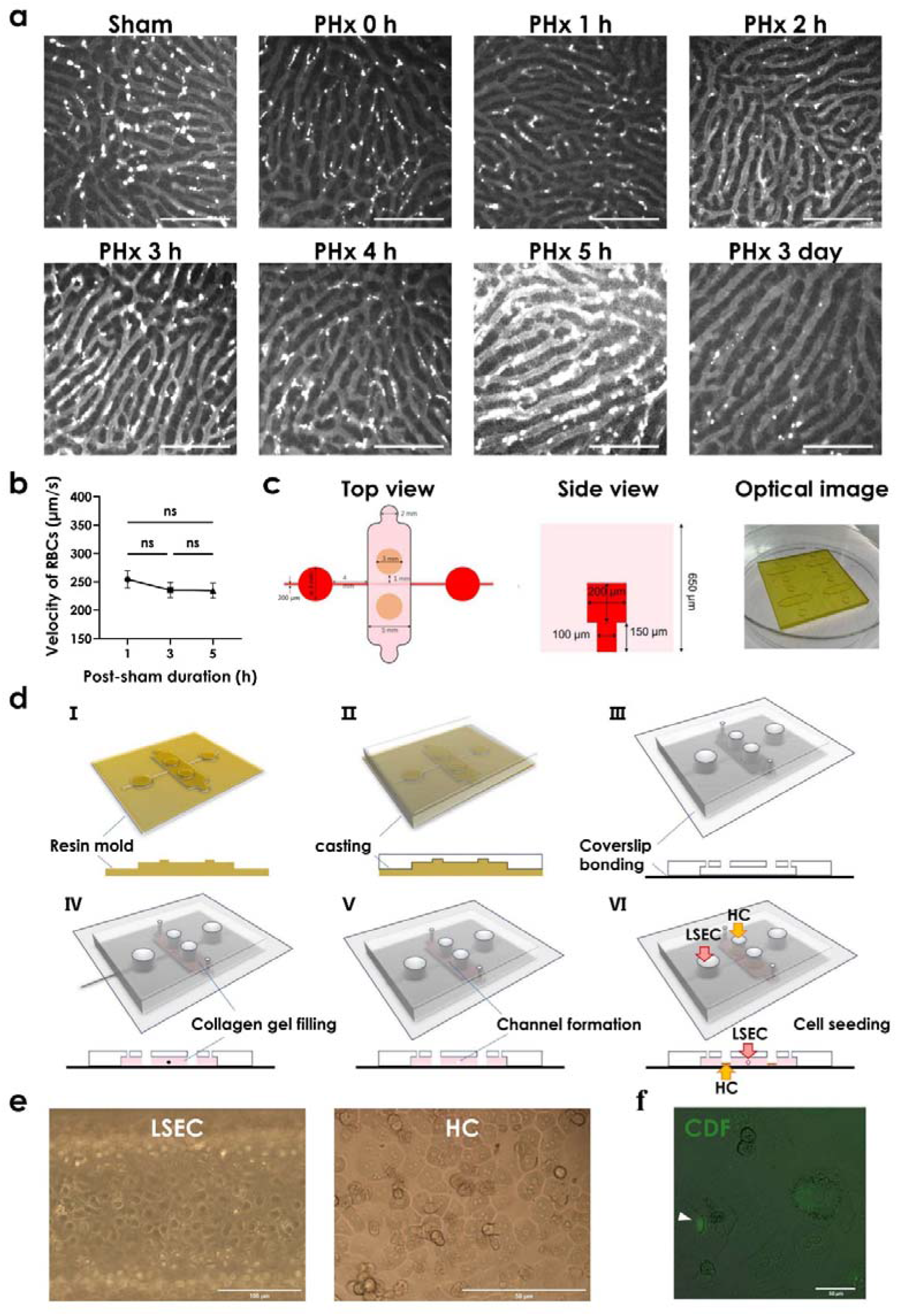
*In vivo* observation in mice and *in vitro* LRC construction. *a*, Representative fluorescent images of hepatic sinusoids at various time points post-hepatectomy, scale bar = 100 μm. *b*, Velocity of moving red blood cells (RBCs) in sinusoids observed at various time points post-sham surgery. Data were presented as the mean ± SEM, *n* = 3-4 independent repeats. *c*, Overview of LRC design. *d*, Fabrication procedure of LRC. *e*, Bright field images of primary LSEC and HC monolayers in LRC. *f*, Fluorescent images of bile canaliculi (*arrow*) between HCs marked stained by CDF.

**Supplementary figure 2.**
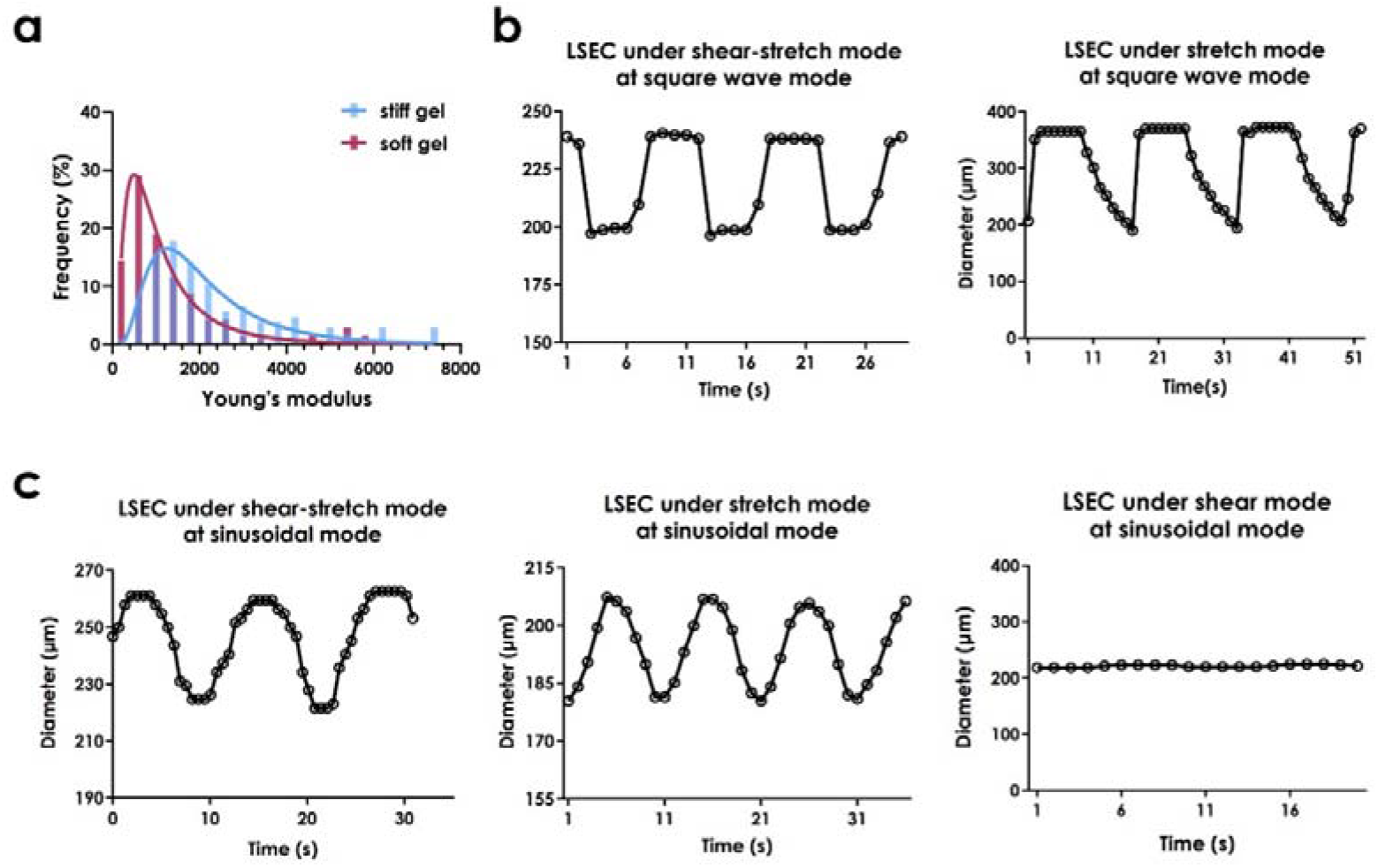
**Characterizing mechanical properties of LRC lumen**. *a*, Young’s modulus of collagen gels at 2.5 (*soft*) and 7.8 mg/ml (*stiff*). Data were collected using AFM assay and represented by typical distributions at a cutoff of < 8000 Pa (yielding the majority of data set > 90%). *b*, Periodically-varied LSEC lumen diameter in LRC with square wave loading under stretch-shear (flow rate at 47 μl/min) and stretch (pressure at 9 mbar) modes. c, Periodically-varied LSEC lumen diameter in LRC with sinusoidal loading under stretch-shear (flow rate at 9.4 μl/min), stretch (pressure at 0.4 mbar) and shear (flow rate at 9.4 μl/min) modes.

**Supplementary figure 3.**
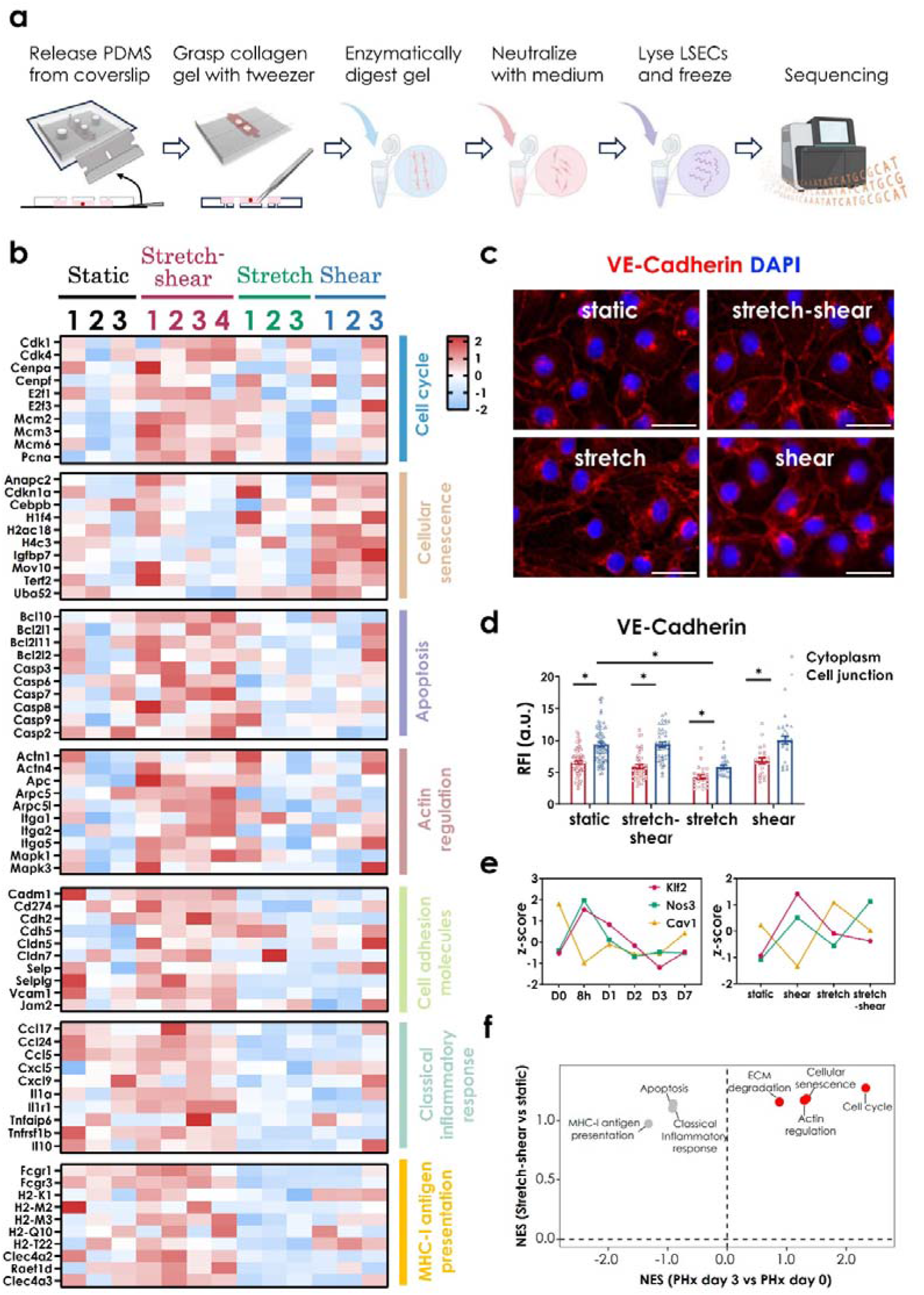
**Sample collection *in vitro* and data comparisons for 3-d post-hepatectomy *in vivo***. *a*, Protocols for collecting RNA samples from LSEC lumen in LRC. *b*, GSEA corresponding heatmap of pathways associated with LSEC phenotype and function under three modes. *c*, *d*, Representative fluorescence images of VE-Cadherin (*c*) in LSECs, with the quantified VE-Cadherin (*d*) distribution for cytoplasm and cellular junction. *e*, Gene expression of Klf2-related targets under mechanical loads in this study and in LSECs from regenerating livers. *f*, GSEA analysis of scRNALseq data for 3-d post-PHx *vs.* sham and stretch-shear *vs.* static in LSECs^37^. Data were collected from stretch-shear (flow rate at 9.4 μl/min) and static modes. *n* = 3-4 independent repeats.

**Supplementary figure 4.**
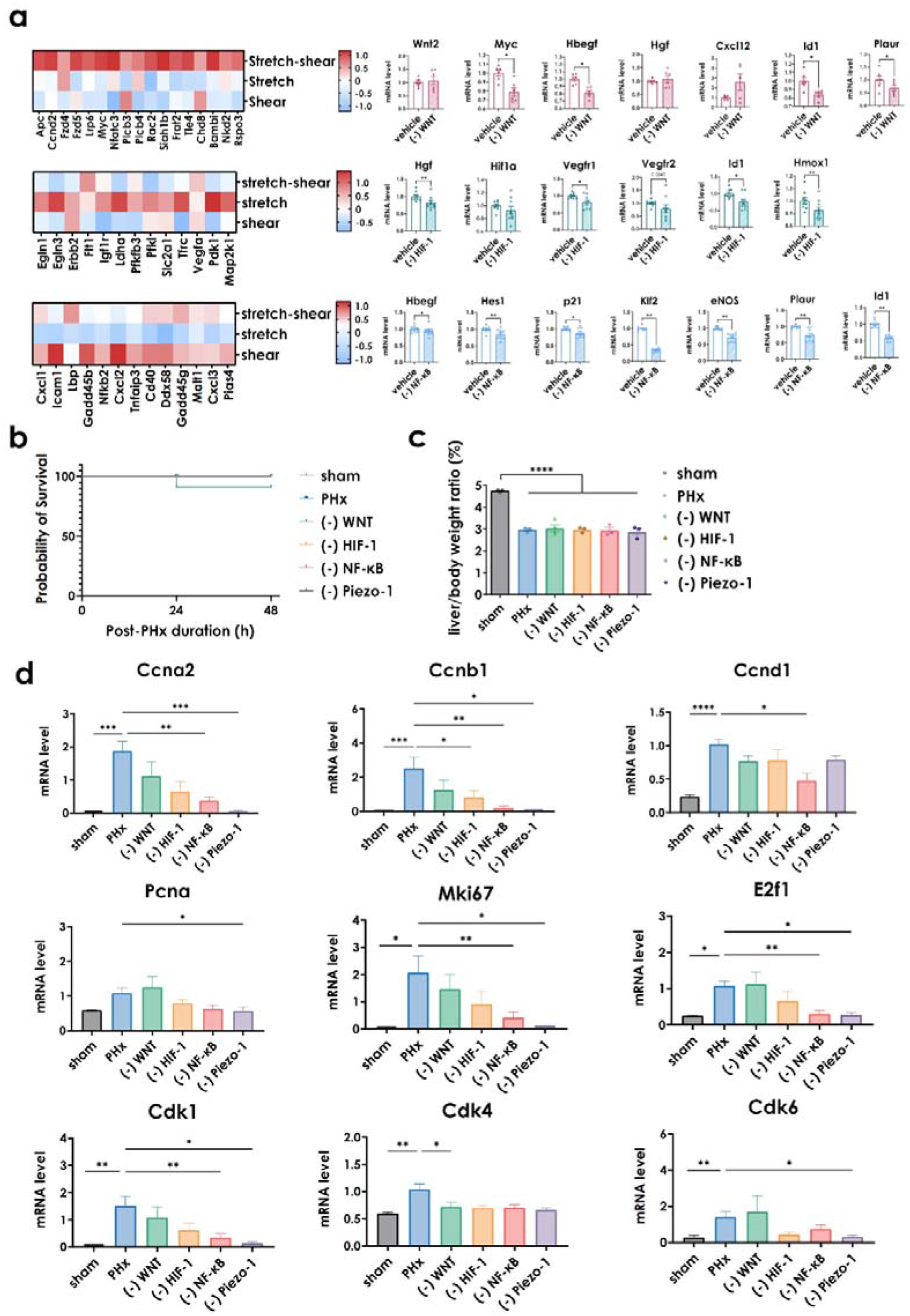
Inhibition of mechano-sensitive pathways for LSECs collected from LRC lumen (*in vitro*) or isolated from PHx mice (*in vivo*). *a*, Heatmap of genes enriched in mechano-sensitive pathways (*left column*) and qPCR test of corresponding inhibitions for Wnt-, HIF-1- and NF-κB-related signaling molecules (*right columns*), respectively. Here IWR-1, LW6, and PDTC denoted as the inhibitor of Wnt, HIF-1 and NF-κB signaling, respectively. *b*, Survival rate in 70% PHx mice with small-molecule or peptide inhibitor treatment. *c*, Liver/Body weight ratio at 48-h post-hepatectomy for mice with inhibition treatment. *d*, qPCR test of proliferation-related genes in liver tissue at 48-h post-hepatectomy with inhibition treatment. Data were collected from mouse livers at 48-h after 70% PHx with twice injections of the inhibitors, and presented as the mean ± SEM, *n* = 3-4 independent repeats.

**Table S1.**
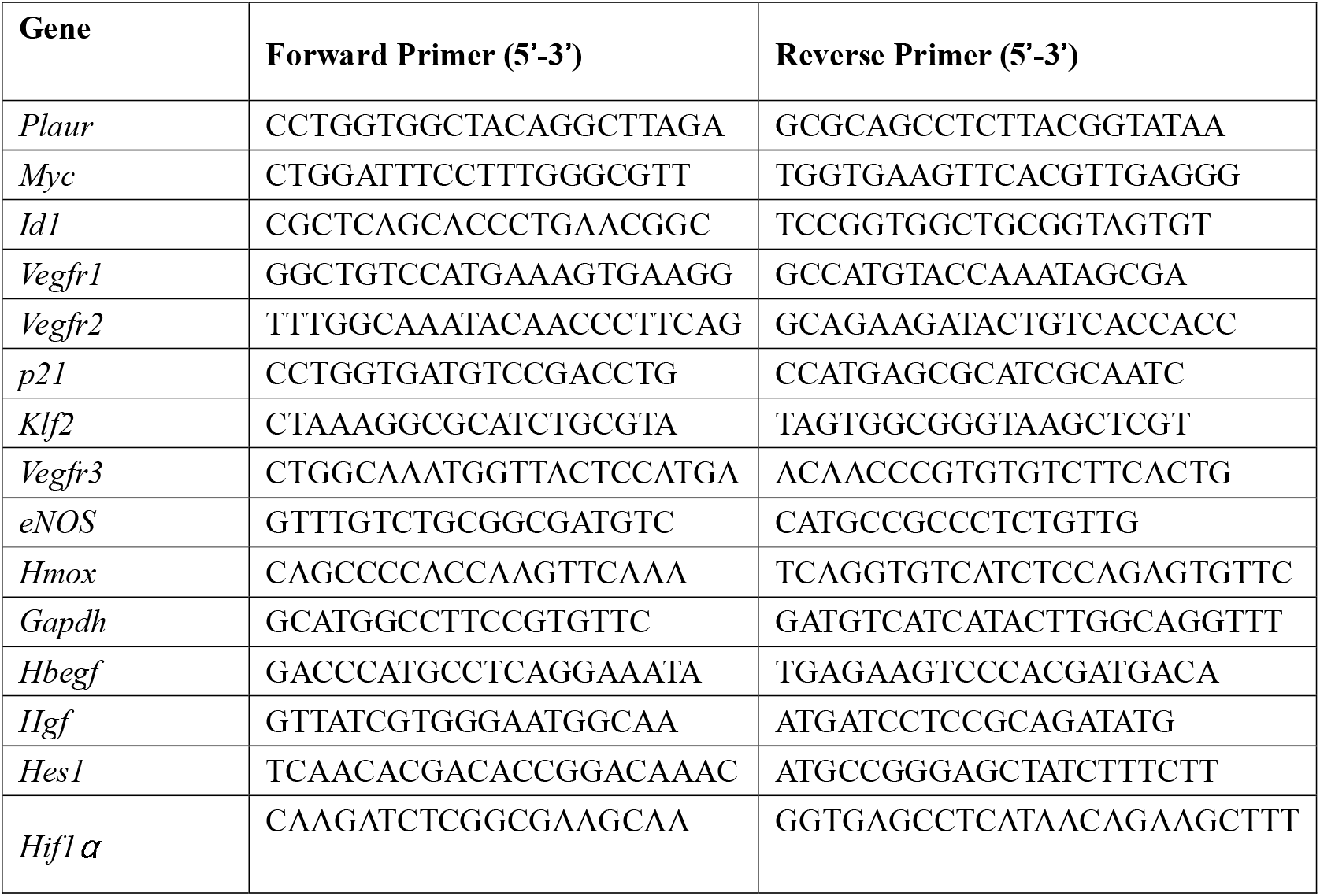
List of PCR primers for qPCR used in this study (related to STAR method)

